# Replacement of tibialis cranialis tendon with polyester, silicone-coated artificial tendon preserves biomechanical function in rabbits

**DOI:** 10.1101/2023.10.25.563771

**Authors:** Katrina L. Easton, Carter Hatch, Kaitlyn Stephens, Dylan Marler, Obinna Fidelis, Xiaocun Sun, Kristin M. Bowers, Caroline Billings, Cheryl B. Greenacre, David E. Anderson, Dustin L. Crouch

**Author notes:** Corresponding author: Department of Mechanical, Aerospace, and Biomedical Engineering University of Tennessee – Knoxville 306D Dougherty Engineering Bldg. 1512 Middle Drive Knoxville, TN 37996. Author Contribution Statement: All authors made one or more of the following contributions to this manuscript: [1] substantial contributions to research design, or the acquisition, analysis or interpretation of data; [2] drafting the paper or revising it critically; [3] approval of the submitted and final versions. All authors have read and approved the final submitted manuscript.

## Abstract

Artificial tendons may be an effective alternative to autologous and allogenic tendon grafts for repairing critically sized tendon defects. The goal of this study was to quantify the in vivo hindlimb biomechanics (ground contact pressure and sagittal-plane motion) during hopping gait of rabbits having a critically sized tendon defect of the tibialis cranialis and either with or without repair using an artificial tendon. In five rabbits, the tibialis cranialis tendon of the left hindlimb was surgically replaced with a polyester, silicone-coated artificial tendon (PET-SI); five operated control rabbits underwent complete surgical excision of the biological tibialis cranialis tendon in the left hindlimb with no replacement (TE). At 8 weeks post-surgery, peak vertical ground contact force in the left hindlimb was statistically significantly less compared to baseline for the TE group (p=0.0215). Statistical parametric mapping (SPM) analysis showed that, compared to baseline, the knee was significantly more extended during stance at 2 weeks post-surgery and during the swing phase of stride at 2 and 8 weeks post-surgery for the TE group (*p*<0.05). Also, the ankle was significantly more plantarflexed during swing at 2 and 8 weeks postoperative for the TE group (p<0.05). In contrast, there were no significant differences in the SPM analysis among timepoints in the PET-SI group for the knee or ankle. These findings suggest that the artificial tibialis cranialis tendon effectively replaced the biomechanical function of the native tendon.

## Introduction

Injuries to tendons and ligaments account for at least 4% of all musculoskeletal trauma cases^1^. Critically-sized tendon defects (i.e., those too large to heal spontaneously), or gaps, are especially debilitating because tendons perform essential biomechanical functions during movement, including storing elastic energy and transmitting forces between muscles and bones. Such defects may form at the time of severe trauma^2,3^ or with chronic tendon ruptures for which the ruptured ends cannot be approximated due to muscle retraction^4^. Autologous tendon grafts represent the current gold-standard and are the most common clinical treatment^5^, although allogenic grafts are also used^6^. Clinical use of autologous grafts is limited by donor site morbidity^7^, while allografts present biosafety concerns and depend on a sufficient supply of donor tendons. Functional outcomes with tendon grafts are modest, with less than half of patients achieving “excellent” function based on standardized clinical assessments^5,8,9^.

Artificial tendons that permanently replace part or all of a biological tendon may be an effective alternative to tendon grafts for critically sized tendon defects. Many types of artificial tendons of varying designs and materials have been tested^10–13^. One recent polyester, silicone-coated (PET-SI) artificial tendon was tested in rabbits^14^ and goats^15–17^ for up to 180 days; the artificial tendon integrated closely with the muscle fibers with no apparent scarring; the muscle-artificial tendon interface was stronger than the muscle itself. Similar artificial tendons have been used in humans^18,19^. However, despite the critical biomechanical role of tendons during movement, the effect of the artificial tendons on movement biomechanics has not been rigorously quantified. We recently reported the hindlimb biomechanics of rabbits with surgical replacement of either the tibialis cranialis or Achilles biological tendon with a PET-SI artificial tendon^20^. For both groups, ankle kinematics and vertical ground contact forces during the stance phase of hopping gait recovered from 2-6 weeks postoperative toward those measured pre-surgery. Three key limitations of this previous study, which motivated the study presented herein, were small treatment groups (n=2), measurements limited to hindlimb biomechanics during the stance phase of hopping gait, and failure of the Achilles artificial tendon at the point of attachment of the artificial tendon to the bone anchor prior to the study endpoint.

The present study was performed to focus on the effect of critically sized defects of the tibialis cranialis tendon on functional mechanics of the hind limb. Our objective was to quantify unilateral hindlimb biomechanics during the entire hopping gait cycle of rabbits having loss of the tibialis cranialis tendon as compared with rabbits having the defect repaired with an artificial tibialis cranialis tendon. Since the tibialis cranialis is an ankle dorsiflexor muscle, our *a priori* hypotheses were that compared to rabbits with tibialis cranialis tendon excision, rabbits with the artificial tendon would have greater (1) maximum ankle dorsiflexion angle and (2) ankle range of motion during the swing phase of gait.

## Methods

### Artificial Tendon

The artificial tendon, adapted from a previously reported design^14–17^, consisted of two custom double-armed strands of size 0 braided polyester suture (RK Manufacturing, Danbury, CT, USA) (Figure 1). The strands were folded in half and braided for the desired length of the artificial tendon; tendons of varying lengths were fabricated in 2 to 4 mm increments to accommodate variation in lengths of the biological tendons they replaced. The folded distal end formed a loop to facilitate attachment to bone with a suture anchor; the proximal end had swaged needles on each strand for suturing the tendon to muscle. The braid was coated with biocompatible silicone (LSR BIO M340, Elkem, Oslo, Norway) to discourage tissue adhesion.

**Figure 1:**
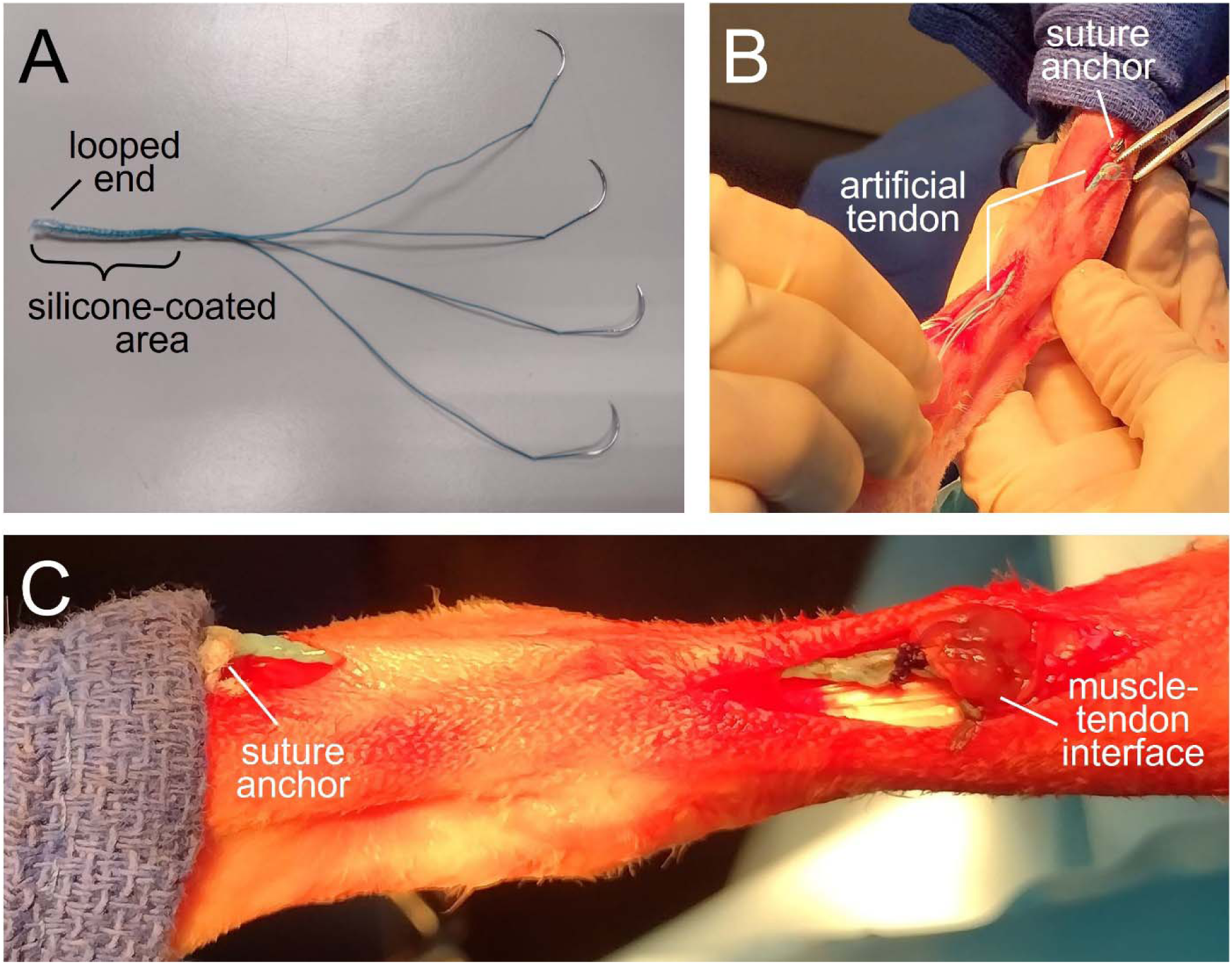
Polyester, silicone-coated (PET-SI) artificial tendon. A) Artificial tendon prior to implantation. The looped end was tied to a suture anchor, while the needle ends were sewn to the distal end of the tibialis cranialis muscle. B) Intraoperative placement of the artificial tendon using two separate skin incisions. C) Completed implantation of artificial tendon prior to closure of incisions.

### Tendon Replacement Model

All animal procedures were approved by the Institutional Animal Care and Use Committee at the University of Tennessee-Knoxville (protocol #2726). All procedures were performed at the University of Tennessee-Knoxville. The study included ten healthy female New Zealand White rabbits (Robinson Services Inc, USA), ranging from 17 to19 weeks old weighing an average of 3.64 ± 0.31 kg (Table 1). Rabbits were housed individually, acclimatized for a minimum of 2 weeks prior to surgery, fed ad libitum (standard laboratory diet, Timothy hay, daily greens), and given daily positive human interaction and enrichment. In addition, rabbits received playpen time twice weekly for at least 10 minutes prior to surgery and starting 2 weeks post-surgery.

**Table 1:** Rabbit demographics and biological and artificial tendon lengths for the tendon excision only and tendon excision with replacement groups.

|  | Age (wks) | Weight at surgery (kg) | Weight at euthanasia (kg) | Biological tendon length (mm) | Artificial tendon length (mm) | Percent Biological Tendon (%) |
| --- | --- | --- | --- | --- | --- | --- |
| **Tendon Excision Only Group** |  |  |  |  |  |  |
| TE1 | 17.4 | 3.08 | 3.9 | 62 | NA | NA |
| TE2 | 17.4 | 3.62 | 4.63 | 60 | NA | NA |
| TE3 | 17.4 | 3.74 | 4.74 | 58 | NA | NA |
| TE4 | 19.4 | 4.02 | 4.87 | 43 | NA | NA |
| TE5 | 19.4 | 3.85 | 4.61 | 51 | NA | NA |
| **Mean (std)** | **18.2 (1.1)** | **3.66 (0.36)** | **4.55 (0.38)** | **54.8 (7.8)** |  |  |
| **Tendon Excision and Replacement Group** |  |  |  |  |  |  |
| PET-SI1 | 19.1 | 4 | 4.65 | 50 | 44 | 88.0 |
| PET-SI2 | 17.4 | 3.69 | 4.36 | 50 | 45 | 90.0 |
| PET-SI3 | 17.4 | 3.21 | 4.26 | 50 | 46 | 92.0 |
| PET-SI4 | 19.4 | 3.75 | 4.56 | 55 | 51 | 92.7 |
| PET-SI5 | 19.4 | 3.42 | 4.47 | 53 | 45 | 84.9 |
| **Mean (std)** | **18.6 (1.0)** | **3.6 (0.31)** | **4.5 (0.16)** | **51.6 (2.3)** | **46.2 (2.8)** | **89.5 (3.2)** |

The rabbits were randomly assigned to either the excision only group (TE, n=5) or the group with replacement of the tendon with a polyester, silicone-coated artificial tendon (PET-SI, n=5). The randomization sequence was generated using a custom script in Matlab (MathWorks, Inc., Natick, MA, USA). To reduce the number of animals, no control group was used; values of variables measured pre-surgery were considered reference or control values. An experimental unit was defined as one rabbit. Potential confounders (e.g., time of surgery, housing location) were not controlled.

The results reported herein are part of a larger overall study that also included measurement of muscle properties (results forthcoming). Thus, the number of animals per group, n=5 (Fig. 5), was computed *a priori* based on a power analysis that considered muscle property values reported in the literature. Specifically, following surgical tenotomy, rabbit soleus muscle mass decreased by 25% compared to intact muscle at four weeks post-tenotomy^21^. We conducted a power analysis (G*Power 3.1, Heinrich-Heine-Universität Düsseldorf, DE) to compute the sample size required to detect *recovery* of 20% over time in the PET group. For a between-timepoint comparison using a two-tailed Student’s t-test, a per-group sample size of n=4 is needed to detect an effect size of 3.46 (i.e. 20% recovery) with power β = 0.90 and significance α = 0.05. We increased the per-group sample size to n=5 to conservatively account for potentially higher within-group variation of muscle mass and other outcome measures.

Rabbits were given hydromorphone (0.1 mg/kg IM) as a preoperative analgesic, sedated with midazolam (1 mg/kg IM), and induced into general anesthesia with isoflurane via face mask. Rabbits were intubated and positioned in right lateral dorsal oblique recumbency. General anesthesia was maintained with isoflurane gas vaporized in 100% oxygen. A loading dose of lidocaine (2 mg/kg IV) was given, followed by a lidocaine CRI (50 mcg/kg/min IV) with isotonic fluids at a rate of 30 ml/h IV throughout the procedure. The left hind limb was clipped, suspended, and aseptically prepared for surgery. A second dose of hydromorphone (0.05 mg/kg IM) was given just prior to the start of surgery.

In the TE group,□1 cm and 2 cm incisions were made over the point of insertion and musculotendinous junction, respectively, of the tibialis cranialis muscle. The tendon was excised from the enthesis to the musculotendinous junction.

In the PET-SI group, a□2 cm incision was made over the point of insertion of the tibialis cranialis muscle. The tendon was released at its insertion. A guide hole was□pre-drilled with a□1.5□mm drill bit□in the proximal metatarsus at the insertion point. A□2 mm x 6 mm□bone suture anchor (Jorgenson Laboratories, Loveland, CO, USA) was screwed into the hole until finger tight. A 3 cm incision was made over the musculotendinous junction of the tibialis cranialis muscle. From among the available artificial tendon lengths, we selected a length that was approximately 85 - 95% of the length of each rabbit’s biological tendon (Table 1) as measured when the foot was in full plantarflexion; the shorter length was selected to offset the length added by the bone suture anchor. The artificial tendon was passed underneath the skin from the proximal incision to the distal incision. The distal loop of the artificial tendon was sutured to the anchor using□#2□Fiberwire□suture with□two□passes□through the anchor and loop. The needle ends of the□artificial tendon were sutured to the distal end of the tibialis□cranialis□muscle using a single suture loop pattern for each strand. Adjacent suture strands were tied together using□six throws, and excess suture was cut and removed. The tibialis□cranialis□tendon was□sharply excised at the musculotendinous junction.□

In both groups, the incisions were closed with a double-layer closure. The subcutaneous layer was closed with a simple continuous pattern with□3-0 PDS□(synthetic absorbable monofilament). The skin was closed□with□an intradermal pattern using□3-0 PDS. An incisional block with 0.15 ml of lidocaine (2 mg/ml SC) was performed. The rabbits received meloxicam (1 mg/kg SC) and enrofloxacin (5 mg/kg SC diluted in 6 ml of sterile saline) immediately post-surgery. As part of a standard protocol for management of post-operative pain and inflammation, all rabbits had laser therapy (MultiRadiance ACTIVet Pro, Solon, OH, USA; 1000 Hz for 1 min, 50 Hz for 1 min, and 1000-3000 Hz for 1 min) performed, once, immediately post-surgery. Left lateral and craniocaudal radiographic views were acquired immediately post-operatively and then every 2 weeks post-surgery.

The operated limb was bandaged for three days post-surgery, and silver sulfadiazine topical cream was applied to the incisions. Each rabbit received hydromorphone (1 mg/kg IM; q 6 h for 3 days), enrofloxacin (5 mg/kg PO; q 12 h for 7 days), and meloxicam (1 mg/kg PO; q 24 h for 7 days). Lactated ringer’s solution (150 ml SC) was administered twice daily starting the day after surgery and continuing for 5 doses. Rabbits were weighed at least twice per week for two weeks post-surgery, then at least every other week for the remainder of the study. Our IACUC protocol established *a priori* that a rabbit would be removed from the study by humane euthanasia if (1) its body weight decreased by at least 20% from the pre-surgery and showed no signs of improvement with intervention; (2) there was dehiscence, tissue breakdown, or infection that could not be treated or repaired; or (3) other signs of distress were present that could not be managed.

Prior to surgery and after recovery from surgery, biomechanics and imaging data were collected every other week until the end of the study. During off weeks, in a single session, the rabbits hopped along the walkway six times in each direction (see below). At 8 weeks post-surgery, the rabbits were humanely euthanized by intravenous overdose of pentobarbital (390 mg/ml, minimum 1 ml/10 lbs).

### Biomechanics Testing

Prior to surgery, rabbits were trained to hop along a 2.6 m-long walkway with an active high-resolution pressure sensing area of 1.3 m (2-Tile High-Resolution Strideway System, Tekscan, Norwood, MA, USA). Reflective 7.5 mm flat circular markers were placed on the lateral aspect of the left limb at the hip (greater trochanter), knee, ankle (lateral malleolus), and 5^th^ metatarsophalangeal (MTP) joint (Figure 2). Marker trajectories were recorded with three high-speed cameras (Prime 13, OptiTrack, NaturalPoint, Inc, Corvallis, OR, USA) that were placed equidistant from each other and parallel to one side of the walkway in order to capture sagittal plane motion. Pressure and video data were recorded synchronously at 240 Hz. The researchers were not blinded to rabbit group assignment during the biomechanics test sessions.

**Figure 2:**
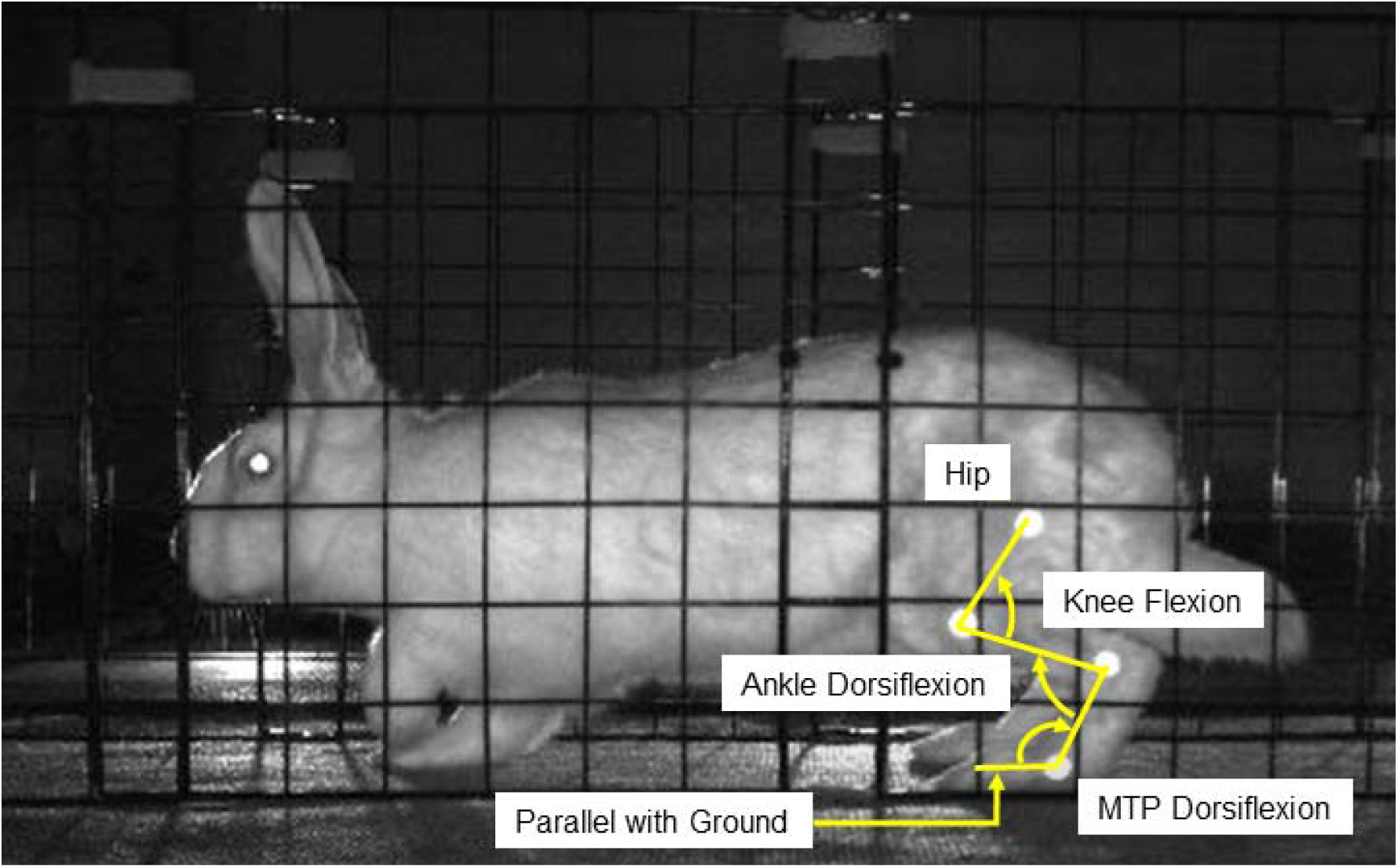
Image of a subject hopping in the walkway. Bright dots are the 7.5 mm flat circular markers located at the hip, knee, ankle, and 5^th^ metatarsophalangeal (MTP) joints. Arrows indicate direction for flexion of a given joint.

Pressure data were processed using pressure analysis software (Strideway 7.80, Tekscan, Norwood, MA, USA) and custom written MATLAB scripts (MATLAB 2022a, MathWorks, Natick, MA, USA). Videos were exported from the motion capture software (Motive:Tracker 1.9, OptiTrack, NaturalPoint, Inc. Corvallis, OR, USA) and marker position data was initially obtained using DeepLabCut^22^. Custom written MATLAB scripts were used to manually verify and adjust the marker positions, calculate sagittal plane joint angles for the knee, ankle, and MTP joints, and combine the joint angle data from the 3 cameras into one time-series curve for the entire length of the walkway. MTP angle during stance was calculated using the ankle marker and a ground plane defined as a horizontal line on the video frames (Figure 2). The researchers who processed the data were not involved in the surgery or data collection, and were blinded to rabbit group assignment.

### Data Analysis

Statistical analysis software (SAS 9.4, SAS Institute, Cary, NC, USA) was used to perform a two-factor (group, timepoint, and group*timepoint) analysis of variance (ANOVA) with repeated measures (rabbit). A *p*-value < 0.05 was used to determine any significant differences. The factor “group” included levels “TE” and “PET-SI”, and the factor “timepoint” included levels “baseline” (pre-surgery), “2 weeks post-surgery”, and 8 weeks post-surgery”. Normality of data was assessed using a Shapiro-Wilk test, and non-normal data were corrected using rank data transformation if necessary. Gait velocity was included as a covariate in the model. Least squared means with a Tukey-Kramer adjustment was used for post-hoc pairwise comparisons, ten in total:

- Weeks Post-Surgery

○ Baseline vs. 2 weeks post-surgery
○ Baseline vs. 8 weeks post-surgery
○ 2 weeks post-surgery vs. 8 weeks post-surgery
- Group x Weeks Post-Surgery

○ TE baseline vs PET-SI baseline
○ TE baseline vs. TE 2 weeks post-surgery
○ TE baseline vs. TE 8 weeks post-surgery
○ TE 2 weeks post-surgery vs. TE 8 weeks post-surgery
○ PET-SI baseline vs. PET-SI 2 weeks post-surgery
○ PET-SI baseline vs. PET-SI 8 weeks post-surgery
○ PET-SI 2 weeks post-surgery vs. PET-SI 8 weeks post-surgery

Three trials for each timepoint for each rabbit were selected to include in the analysis (3 trials x 3 timepoints x 10 rabbits = 90 trials). In order to choose the three trials, each video initially was assessed qualitatively; a video was excluded if the rabbit was hopping abnormally (i.e., play hopping or flicking of feet while hopping) at any point during the trial. For the remaining videos, the gait velocity was calculated for each rabbit and video and averaged across timepoints. At each timepoint, the three trials that were closest to the average gait velocity for the rabbit were selected for further analysis. Comparisons were made for the operated limb only. The independent variables for pressure mat data were peak vertical force, vertical impulse, vertical impulse distribution, and average ground contact area. The independent variables for kinematics data were stance percent of stride, range of motion, and maximum, minimum, and average joint angle for the knee and ankle during the stance and swing phase of gait and for the MTP during stance phase of gait. Stance and swing phases of gait were determined using the pressure mat data. Peak vertical force and vertical impulse were normalized by body weight.

Statistical parametric mapping (SPM)^23,24^ open-source MATLAB software^25^ was used to compare the kinematic curves for each group, joint, and gait phase. A two-factor (group, timepoint, and group*timepoint) ANOVA with repeated measures (rabbit) was performed first. Then, post-hoc tests were performed using a SPM two-tailed paired t-test to compare joint angles between timepoints within each group. A Bonferroni correction was used to account for multiple comparisons.

## Results

No rabbit was excluded or removed prematurely from the study; all rabbits in each group (n=5 per group) were included in the data analysis. There were no significant differences in age or weight at time of surgery, weight at euthanasia, or length of the biological tendon between the two groups (Table 1). The length of the artificial tendons ranged from 84.9% to 92.7% (mean 89.5 ± 3.2%) of the length of the biological tendon (Table 1), which was within the targeted range.

### Biomechanics: Pressure Data

Timepoint was significant for peak vertical force (p=0.0002) and vertical impulse (p=0.0180). More specifically, across groups, both peak vertical force and vertical impulse were significantly less at 2 and 8 weeks post-surgery compared to baseline (Figure 3A and C). In addition, peak vertical force was significantly less at 8 weeks post-surgery compared to baseline for the TE group (*p*=0.0215), but not the PET-SI group (*p*=0.0621). Faster gait velocity was significantly associated with greater peak vertical force (*p*=0.0028) and vertical impulse (*p*=0.0191).

**Figure 3:**
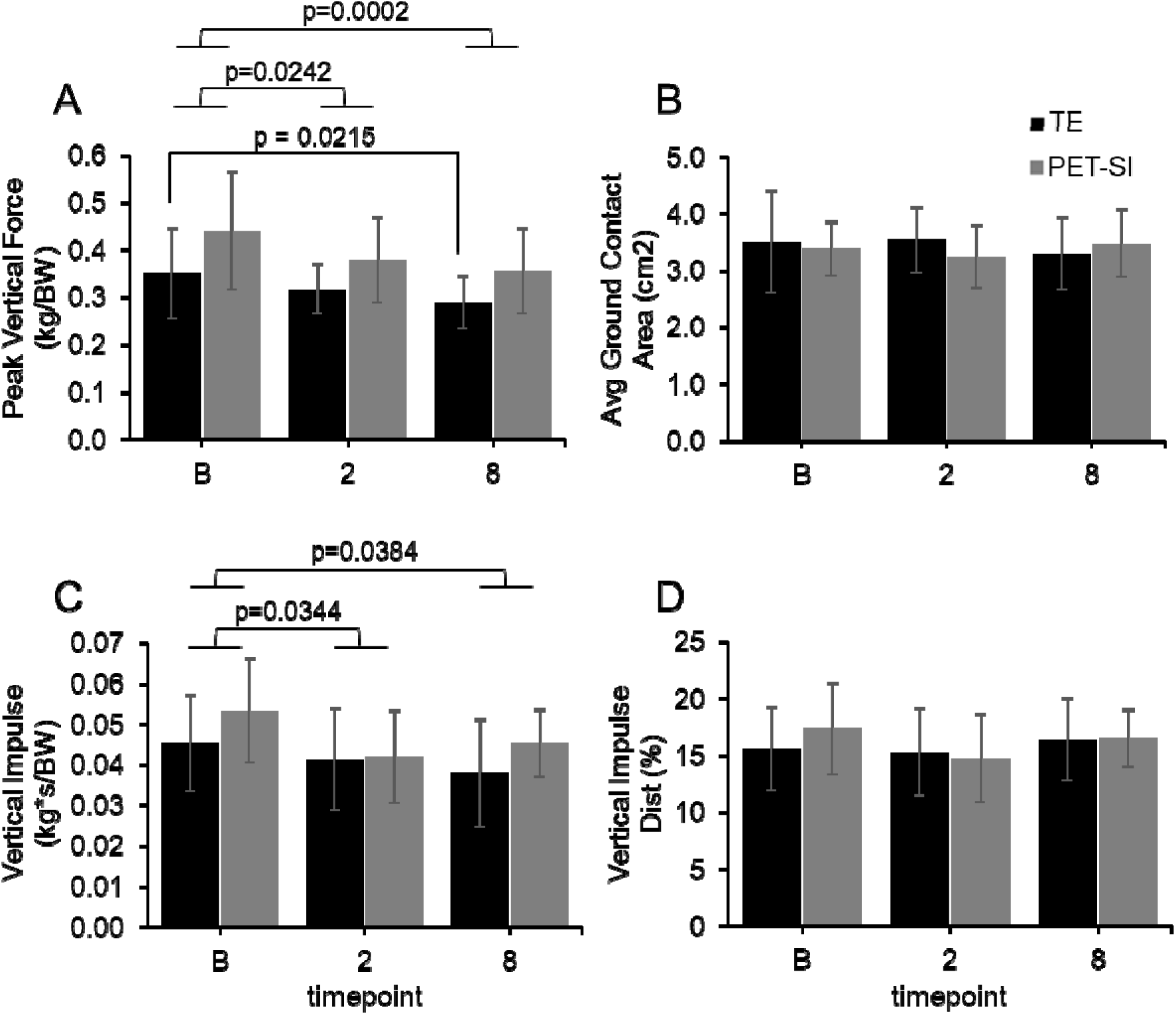
Mean values and standard deviation at baseline (pre-surgery), and 2 weeks, and 8 weeks post-surgery for the operated limb for the tendon excision only (TE) and tendon excision and replacement (PET-SI) groups for A) Peak vertical force normalized by body weight (kg/BW); B) Average ground contact area (cm^2^); C) Impulse normalized by body weight (kg*s/BW); D) Vertical impulse distribution (%).

### Biomechanics: Kinematics

Of the main factors, treatment group was not significant, but timepoint had a significant effect on many of the kinematics variables for both groups (Figures 3 – 5, Table 2). The p-values for main effects, their interaction, and the by-week comparisons are in Tables 2 and 3. Across groups, there was an increase in ankle plantarflexion and knee extension and a decrease in ankle dorsiflexion and knee flexion at 2 and 8 weeks post-surgery compared to baseline, especially during swing phase. Across groups, the overall range of motion during swing phase was significantly less for the knee at 2 weeks post-surgery and the ankle at 2 and 8 weeks post-surgery compared to baseline. There was significant recovery of range of motion in the knee and average angle of the MTP by 8 weeks post-surgery. There was no difference in stance percent of stride across groups or timepoints (Table 4). However, gait velocity was significantly correlated with the stance percent of stride (*p* < 0.0001; Table 2).

**Table 2:** Statistical results (*p*-values) from the 2-factor ANOVA with repeated measures for each kinematic variable for knee, ankle, and MTP joints during stance and swing phase of gait. The normality column indicates whether the data needed to be transformed or not. Group was TE or PET-SI. Weeks post was baseline, 2 or 8-weeks post-surgery. GxW is the group by weeks post-surgery interaction term. Gait velocity was included as a covariate in the model. ROM – range of motion; NS – not significant. *p* < 0.05 were considered significant differences. *p* < 0.1 were also included.

| Knee |  |  |  |  |  |  |
| --- | --- | --- | --- | --- | --- | --- |
| Variable | Gait Phase | Normality | Group | Weeks Post | GxW | Gait Velocity |
| Max Angle | Stance | NS | NS | 0.027 | NS | NS |
|  | Swing | NS | NS | < 0.0001 | 0.0008 | NS |
| Min Angle | Stance | Transformed | NS | 0.0932 | NS | NS |
|  | Swing | NS | NS | < 0.0001 | 0.0108 | NS |
| Avg Angle | Stance | Transformed | NS | 0.0809 | 0.0497 | NS |
|  | Swing | Transformed | NS | 0.0004 | 0.0066 | NS |
| ROM | Stance | NS | NS | NS | NS | NS |
|  | Swing | NS | NS | 0.0017 | NS | NS |
| Ankle |  |  |  |  |  |  |
| Variable | Gait Phase | Normality | Group | Weeks Post | GxW | Gait Velocity |
| Max Angle | Stance | Transformed | NS | NS | NS | 0.0575 |
|  | Swing | Transformed | NS | 0.0145 | NS | NS |
| Min Angle | Stance | NS | NS | 0.0012 | NS | NS |
|  | Swing | Transformed | NS | < 0.0001 | NS | NS |
| Avg Angle | Stance | Transformed | NS | 0.0097 | NS | NS |
|  | Swing | Transformed | NS | < 0.0001 | NS | NS |
| ROM | Stance | NS | NS | 0.0825 | NS | NS |
|  | Swing | NS | NS | 0.0055 | NS | NS |
| MTP |  |  |  |  |  |  |
| Variable | Gait Phase | Normality | Group | Weeks Post | GxW | Gait Velocity |
| Max Angle | Stance | NS | NS | 0.0005 | NS | NS |
|  | Swing | Transformed | NS | < 0.0001 | NS | NS |
| Min Angle | Stance | NS | NS | NS | NS | NS |
|  | Swing | NS | NS | 0.0588 | NS | NS |
| Avg Angle | Stance | NS | NS | 0.0398 | NS | NS |
|  | Swing | NS | NS | < 0.0001 | NS | NS |
| ROM | Stance | NS | NS | 0.08 | NS | NS |
|  | Swing | Transformed | NS | NS | NS | NS |

**Table 3:**
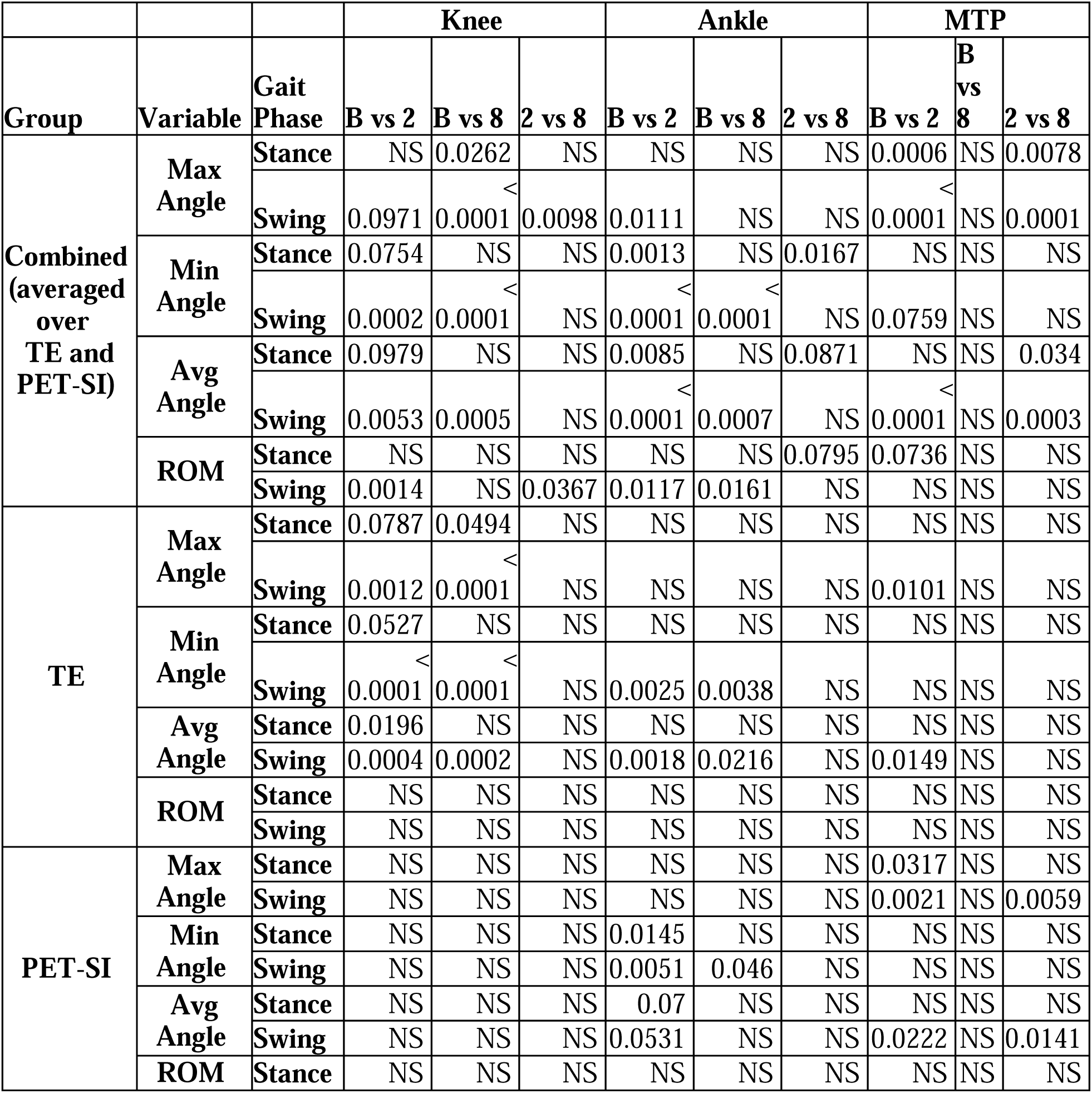
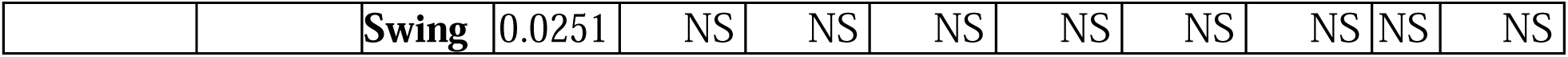
Statistical results (*p*-values) for the LS means with Tukey-Kramer multiple comparison adjustment for each kinematic variable for the knee, ankle, and MTP joints during stance and swing phase of gait for each timepoint. Group was combined (averaged over TE and PET-SI), TE or PET-SI. Timepoints were baseline (B), 2-, or 8-weeks post-surgery. ROM – range of motion; NS – not significant. *p* < 0.05 were considered significant differences. *p* < 0.1 were also included.

**Table 4:** The percentage of the gait cycle that was the stance phase for both groups over time. There were no significant differences between groups or over time.

| **Weeks Post-Surgery/**<br>**Group** | **Stance Percent Stride (Std)** |  |  |
| --- | --- | --- | --- |
|  | **Baseline** | **2** | **8** |
| **TE** | 48.1 (6.0) | 47.6 (4.7) | 48.2 (4.9) |
| **PET-SI** | 48.6 (3.5) | 45.9 (6.9) | 49.7 (5.0) |

For the group-by-timepoint interactions, during stance (Figures 3a, 4a, and 5a, Table 3), the TE group’s knee extension significantly increased from baseline to 8 weeks post-surgery, and the knee was significantly more extended overall at 2 weeks post-surgery compared to baseline. In contrast, the PET-SI group had no significant differences among timepoints for the knee during stance. However, the PET-SI group had significantly less ankle and MTP dorsiflexion at 2 weeks post-surgery compared to baseline.

**Figure 4:**
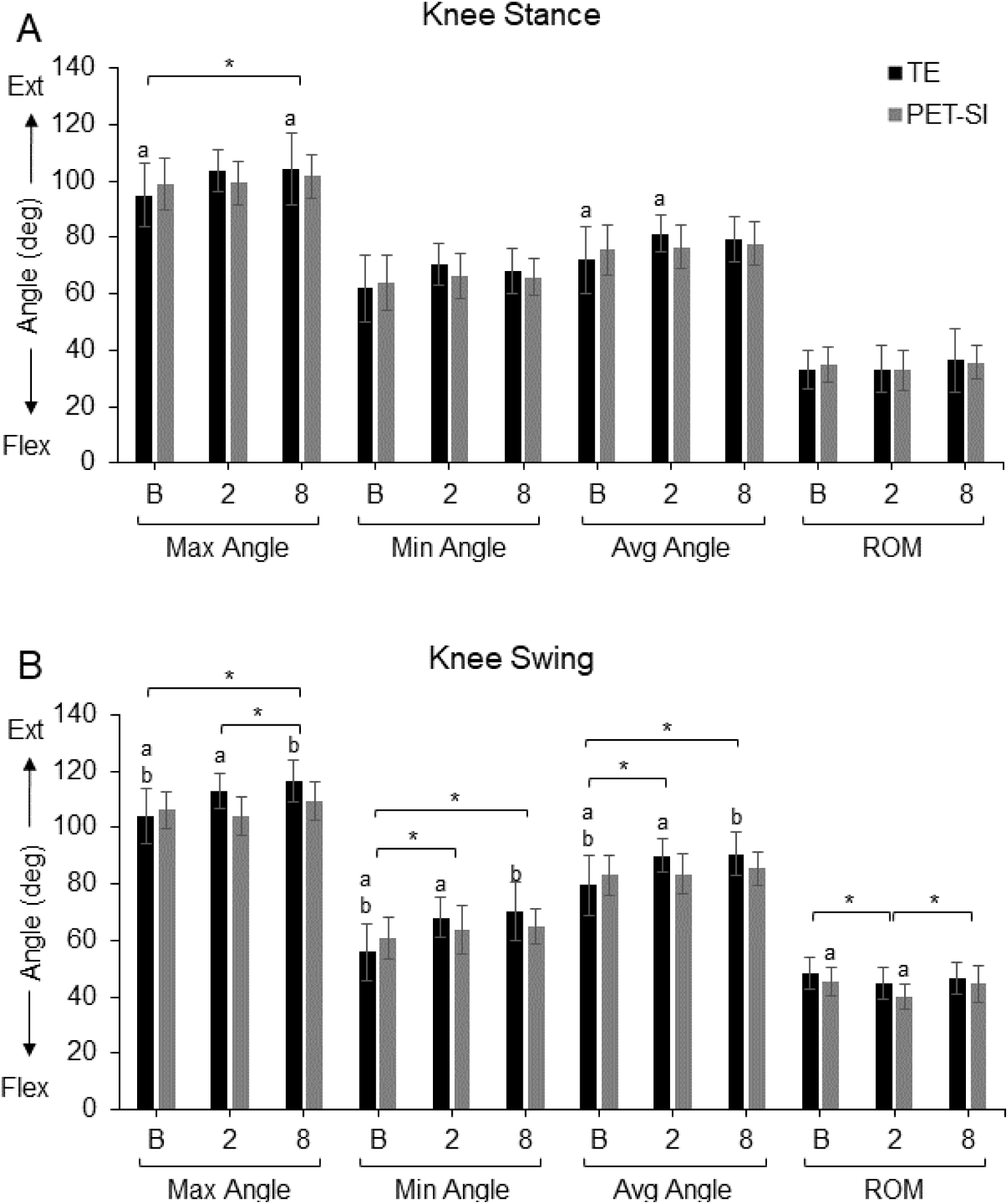
Maximum, minimum, and average knee joint angles and knee range of motion (ROM) for the tendon excision (TE) only group and tendon excision and replacement (PET-SI) group at baseline (B, pre-surgery), 2 weeks post-surgery, and 8 weeks post-surgery during A) stance phase of gait, and B) swing phase of gait. Error bars represent one standard deviation. Ext – Extension; Flex – Flexion. A larger angle indicates greater extension/less flexion. * indicates significant (*p* < 0.05) differences between timepoints across groups. a, b indicate significant (*p* < 0.05) differences between timepoints within a group. Table 3 gives specific *p*-values for each comparison.

There were more between-factor differences in kinematics variables during the swing phase of gait compared to the stance phase (Figures 3b, 4b, 5b, Table 3). At 2 and 8 weeks post-surgery compared to baseline, the TE group maintained the knee and ankle in a significantly more extended/plantarflexed position overall, with significantly greater knee extension, less knee flexion, and less ankle dorsiflexion. For the PET-SI group, the knee range of motion was significantly less at 2 weeks post-surgery compared to baseline, but partly recovered by 8 weeks post-surgery. Similar to the TE group, there was a significant decrease in maximal ankle dorsiflexion at 2 and 8 weeks post-surgery compared to baseline. However, the overall posture (average angle) of the ankle was not significantly affected.

**Figure 5:**
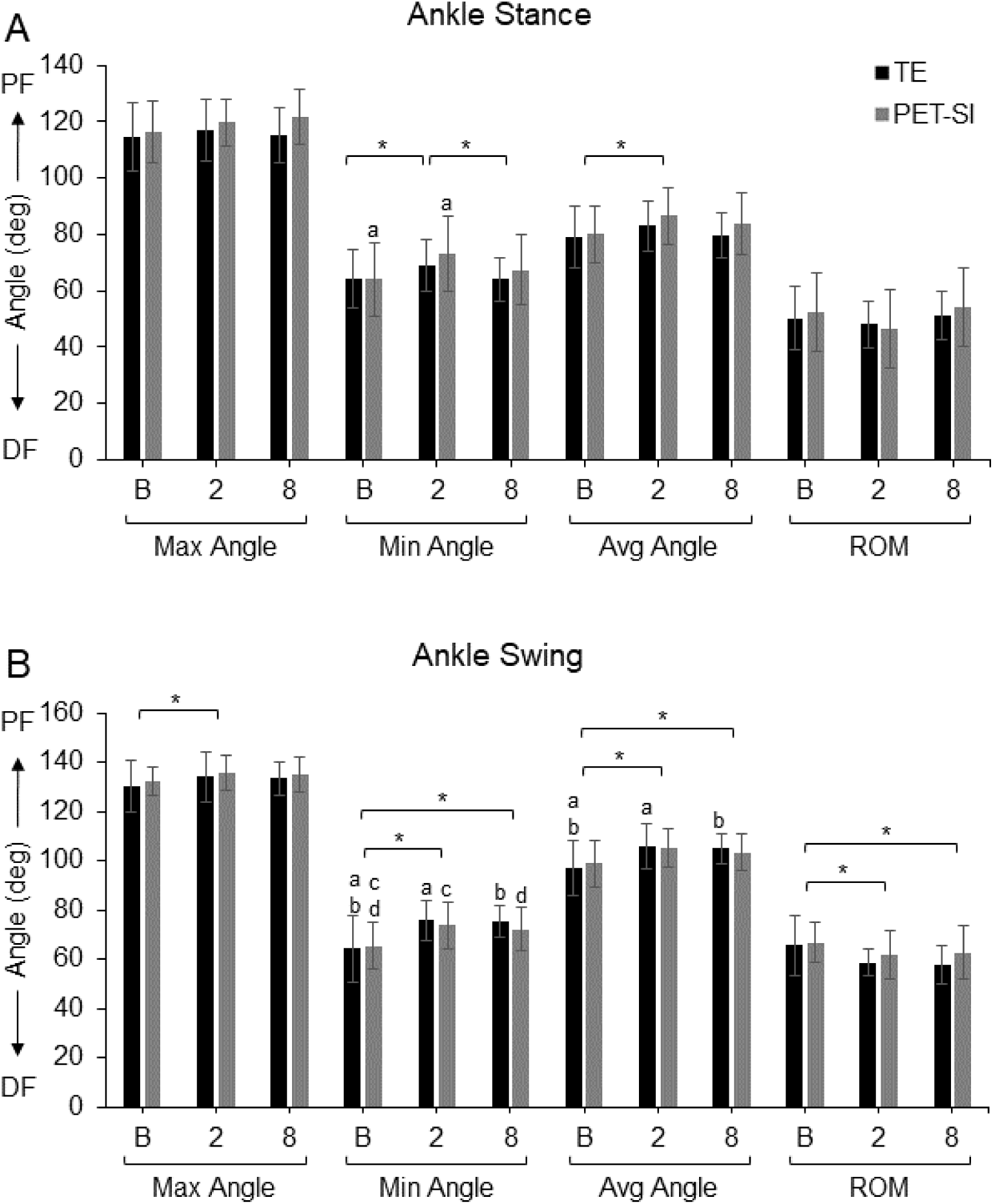
Mean maximum, minimum, and average ankle joint angles and ankle range of motion (ROM) and standard deviation for the tendon excision (TE) only group and tendon excision and replacement (PET-SI) group at baseline (B, pre-surgery), 2 weeks post-surgery, and 8 weeks post-surgery during A) stance phase of gait, and B) swing phase of gait. PF – Plantarflexion; DF – Dorsiflexion. A larger angle indicates greater plantarflexion/less dorsiflexion. * indicates significant (*p* < 0.05) differences between timepoints across groups. a, b indicate significant (*p* < 0.05) differences between timepoints within a group. Table 3 gives specific *p*-values for each comparison.

**Figure 6:**
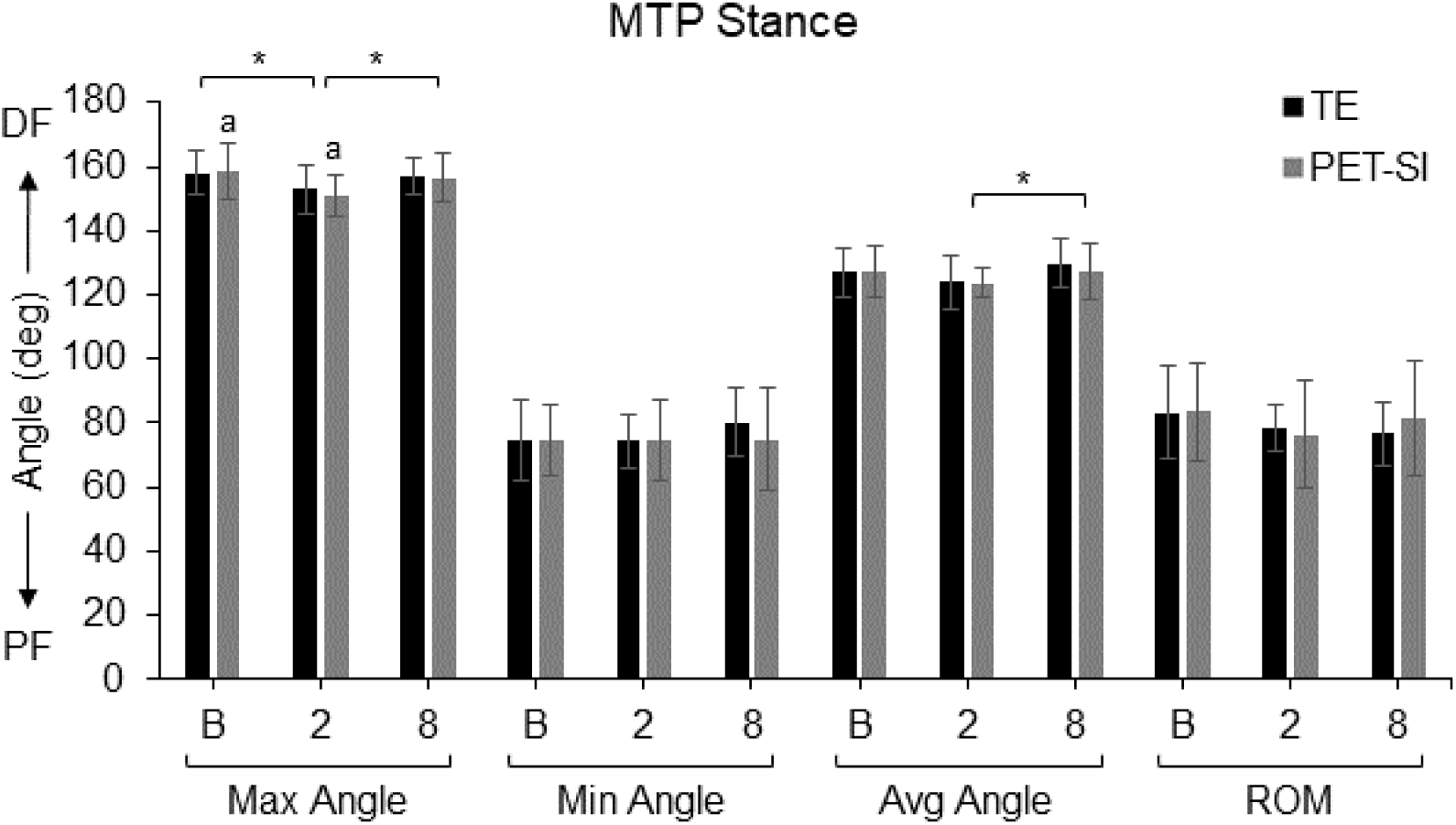
Maximum, minimum, and average MTP joint angles and MTP range of motion (ROM) for the tendon excision (TE) only group and tendon excision and replacement (PET-SI) group at baseline (B, pre-surgery), 2 weeks post-surgery, and 8 weeks post-surgery during stance phase of gait. Error bars represent one standard deviation. PF – Plantarflexion; DF – Dorsiflexion. A larger angle indicates greater dorsiflexion/less plantarflexion. * indicates significant (*p* < 0.05) differences between timepoints across groups. a indicates significant (*p* < 0.05) differences between timepoints within a group. Table 3 gives specific *p*-values for each comparison.

### Statistical Parametric Mapping Analysis

There was a significant difference in joint angle among the three timepoints between ∼44-58% of the swing phase for the knee and ∼38-80% of the swing phase for the ankle. Post-hoc paired t-tests showed that knee flexion was significantly less during stance at 2 weeks post-surgery and during swing at 2 and 8 weeks post-surgery for the TE group (Figure 7a). Ankle dorsiflexion was also significantly less during swing at 2 and 8 weeks post-surgery for the TE group (Figure 7c). There were no significant differences among timepoints in the PET-SI group for the knee or ankle.

**Figure 7:**
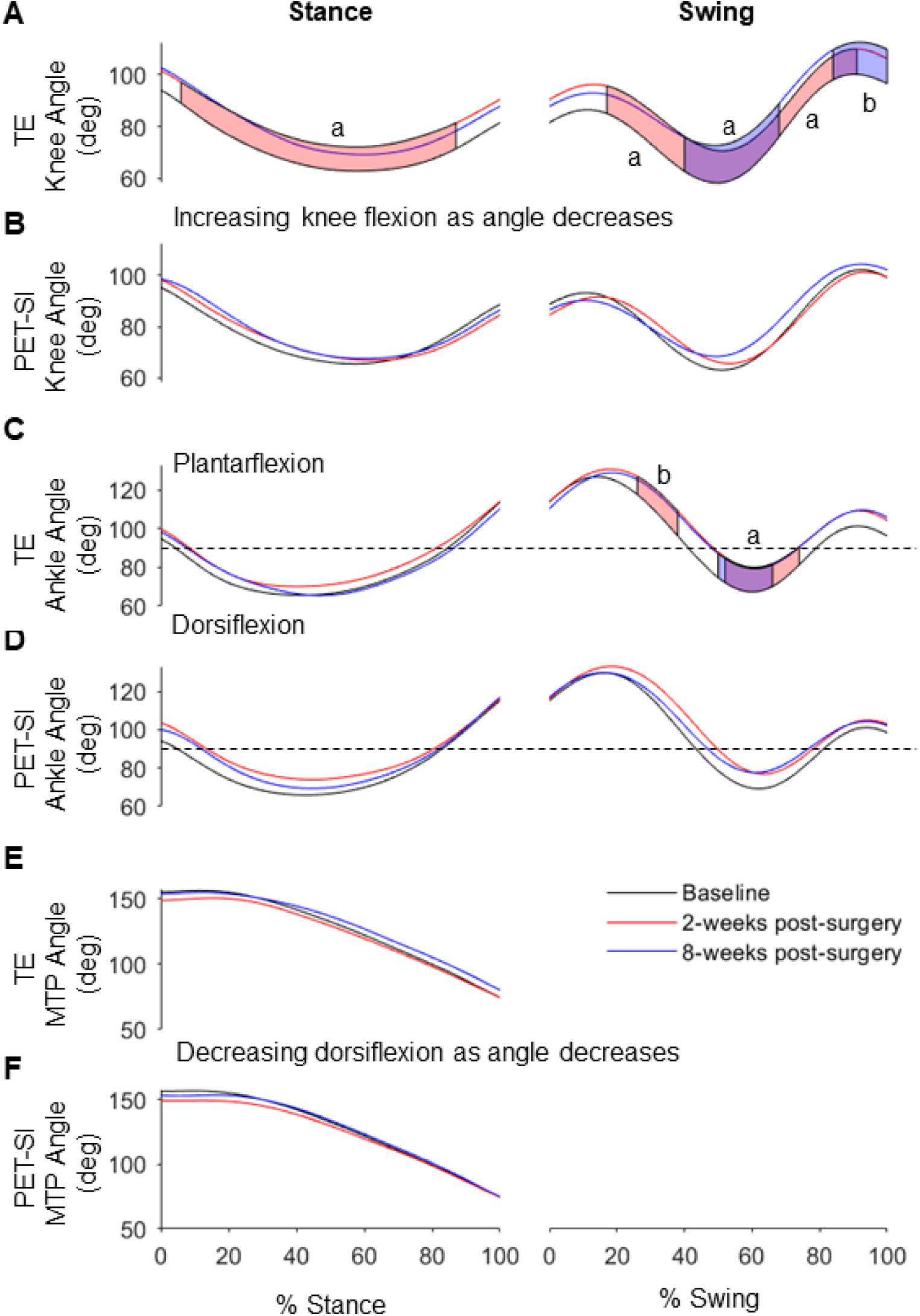
Mean joint angles for TE and PET-SI groups over the gait cycle for the A,B) knee, C,D) ankle, and E,F) MTP. The horizontal dashed line in C and D indicates 90° where the ankle is at neutral dorsi/plantar-flexion. Red shaded areas indicate significant differences between baseline and 2 weeks post-surgery. Blue shaded areas indicate significant differences between baseline and 8 weeks post-surgery. Purple shaded areas are the overlap of red and blue shaded areas. a: *p* = 0.001; b: *p* = 0.002.

## Discussion

The results supported our hypotheses that compared to the TE group, the PET-SI group would have greater peak ankle dorsiflexion angle and ankle range of motion during the swing phase of gait. Additionally, compared to the TE group, the hindlimb kinematics of the PET-SI group were more similar to baseline kinematics measured pre-surgery. Our results were consistent with those of our previous preliminary study^20^ and suggest that the artificial tendons effectively performed the biomechanical function of the native tendons they replaced.

Non-intuitively, peak vertical force at 8 weeks post-surgery was significantly less than at pre-surgery for the TE group and nearly so for the PET-SI group. This was unexpected since the tibialis cranialis muscle, based on its moment arm, does not generate torque that is expected to contribute to vertical ground contact force. A possible explanation for the difference is that the rabbits could have shifted biomechanical loads from the operated limb to the sound limb and spend less time on the operated limb, potentially due to discomfort or perceived functional impairment of the operated limb; further analysis of bilateral pressure data is needed to confirm this. In addition, a longer study duration would help determine if peak vertical force normalizes with additional recovery time.

An important anatomical feature that allows biological tendons to slide relative to surrounding tissues is the tendon sheath. In the PET-SI group, it appeared that a sheath formed around the artificial tendon; immediately post-mortem, we observed the artificial tendon sliding relative to surrounding tissues during passive ankle motion (supplementary video). Thus, the silicone coating performed the intended function of preventing tissues from adhering to the surface of the artificial tendon. These findings are qualitative, visual inspections of the tissues; the extent to which the structure and function of the tissues surrounding the artificial tendons mimic the native epitenon or tendon sheath surrounding the biological tendons is unknown. Future studies will evaluate histology samples to assess the cellular structure of the tissues ensheathing the artificial tendons. Functionally, it would also be useful to quantify the effective mechanical friction and damping coefficients at the tendon-sheath interface.

Though we observed statistically significant differences between the PET-SI and TE groups, the TE group still appeared to have substantial ankle dorsiflexion function. Notably, the TE group had an average maximum dorsiflexion angle during swing phase of gait of 75.4°, which was only 3.1° less than that of the PET-SI group. One possible explanation is that, in the TE group, other muscles crossing the ankle, such as extensor digitorum longus and the peroneus muscle group (longus, brevis, tertius)^26^, may have compensated substantially for the lost contribution of the tibialis cranialis muscle to ankle dorsiflexion torque and movement. The extensor digitorum longus muscle also crosses the knee and ankle^27^ and, thus, may generate an ankle dorsiflexion torque passively as the knee is flexed. Finally, we suspect that the ankle joint is at least partly dorsiflexed passively during both stance and swing phases of hopping gait: during stance as the torso and proximal hindlimb move cranially and near the end of the cranial swing of the hindlimb due to the inertia of the foot.

There are several possible reasons for the modest biomechanical declines from pre- to post-surgery of rabbits with the PET-SI artificial tendons. For one, the PET-SI artificial tendon did not interface with the muscle as seamlessly as a biological tendon does. Specifically, biological tendons integrate with muscles over a large portion of their length via aponeuroses, which facilitates force transmission between muscle and tendon^28^; conversely, a relatively small number (four) of suture strands of the artificial tendon were tied to the distal end of the muscle. Second, the mechanical properties (e.g., stiffness) may be different between biological and artificial tendons; such differences have not been quantified but will be investigated in a future study. Third, though surgical interventions may have caused pain and discomfort in both groups, these may have been greater in the PET-SI group due to the implantation of the artificial tendons. Fourth, the biologic tendon passes under the extensor retinaculum whereas the artificial tendon, due to its size, cannot be placed under the retinaculum. This results in a change in the moment arm and torque direction between the biologic and artificial tendons. Finally, both groups were bandaged for 3 days post-surgery, which partly immobilized the ankle joint; bandaging may have had modest adverse effects (e.g., disuse muscle atrophy) leading to impaired biomechanical function^29^.

There were several limitations of our study. First, the number of samples per group was relatively small, which may have underpowered our statistical comparisons for some variables; however, the group size was similar to those of the previous *in vivo* studies of the PET-SI tendons^14–17^. Second, the study duration was relatively short; functional recovery may have been improved with more time. Third, the study did not include a control group of healthy, non-operated rabbits to account for potential changes in biomechanics as the rabbits aged or gained more experience with our experiment protocol from pre- to post-surgery. Fourth, we surgically replaced a healthy biological tendon with an artificial tendon in a single surgery; future studies should model the common clinical scenario that a tendon defect is present for some period prior to surgical replacement with an artificial tendon.

In conclusion, our *in vivo* study provided the most substantial quantitative evidence to date of a positive treatment effect of PET-SI artificial tendons on biomechanical motor function. Therefore, PET-SI artificial tendons potentially could be an effective alternative treatment option for critically sized tendon defects and other severe tendon pathologies.

## Supporting information

Supplemental Video - Tibialis Cranialis artificial tendon sliding

## Conflict of Interest

None.

## Acknowledgements

Thank you to Dr. Bryce Burton, Dr. Kelsey Finnie, and Chris Carter for veterinary care and support, and to the Animal Housing Facility staff for animal care assistance. Thank you to Kellie McGhee, Ivy Milligan, and Raina Desai for their help with data collection and analysis. Research was funded by the National Institutes of Health Award #R61AR078096. The funder had no role in study design, data collection and analysis, decision to publish, or preparation of the manuscript.

## Notes

### Competing Interest Statement

The authors have declared no competing interest.

## References

1. Clayton RA, Court-Brown CM. The epidemiology of musculoskeletal tendinous and ligamentous injuries. Injury. 2008;39(12):1338–1344.

2. Jepegnanam TS, Nithyananth M, Boopalan PR, Cherian VM, Titus VT. Reconstruction of open contaminated achilles tendon injuries with soft tissue loss. J Trauma. 2009;66(3):774–779.

3. Iorio ML, Han KD, Evans KK, Attinger CE. Combined Achilles tendon and soft tissue defects: functional outcomes of free tissue transfers and tendon vascularization. Ann Plast Surg. 2015;74(1):121–125.

4. Schachter AK, White BJ, Namkoong S, Sherman O. Revision reconstruction of a pectoralis major tendon rupture using hamstring autograft: a case report. Am J Sports Med. 2006;34(2):295–298.

5. Sarzaeem MM, Lemraski MM, Safdari F. Chronic Achilles tendon rupture reconstruction using a free semitendinosus tendon graft transfer. Knee Surg Sports Traumatol Arthrosc. 2012;20(7):1386–1391.

6. Crossett LS, Sinha RK, Sechriest VF, Rubash HE. Reconstruction of a ruptured patellar tendon with achilles tendon allograft following total knee arthroplasty. J Bone Joint Surg Am. 2002;84(8):1354–1361.

7. Kartus J, Movin T, Karlsson J. Donor-site morbidity and anterior knee problems after anterior cruciate ligament reconstruction using autografts. Arthroscopy. 2001;17(9):971–980.

8. LaSalle WB, Strickland JW. An evaluation of the two-stage flexor tendon reconstruction technique. J Hand Surg Am. 1983;8(3):263–267.

9. Frakking TG, Depuydt KP, Kon M, Werker PM. Retrospective outcome analysis of staged flexor tendon reconstruction. J Hand Surg Br. 2000;25(2):168–174.

10. Murray GA, Semple JC. A review of work on artificial tendons. J Biomed Eng. 1979;1(3):177–184.

11. Rawlins R. The role of carbon fibre as a flexor tendon substitute. Hand. 1983;15(2):145–148.

12. Howard CB, McKibbin B, Ralis ZA. The use of Dexon as a replacement for the calcaneal tendon in sheep. J Bone Joint Surg Br. 1985;67(2):313–316.

13. Mendes DG, Iusim M, Angel D, Rotem A, Mordehovich D, Roffman M, Lieberson S, Boss J. Ligament and tendon substitution with composite carbon fiber strands. J Biomed Mater Res. 1986;20(6):699–708.

14. Melvin DB, Klosterman B, Gramza BR, Byrne MT, Weisbrode SL, Litsky AS, Clarson SJ. A durable load bearing muscle to prosthetic coupling. ASAIO journal (American Society for Artificial Internal Organs : 1992). 2003;49(3):314–319.

15. Melvin AJ, Litsky AS, Mayerson JL, Stringer K, Juncosa-Melvin N. Extended healing validation of an artificial tendon to connect the quadriceps muscle to the Tibia: 180-day study. J Orthop Res. 2012;30(7):1112–1117.

16. Melvin A, Litsky A, Mayerson J, Stringer K, Melvin D, Juncosa-Melvin N. An artificial tendon to connect the quadriceps muscle to the tibia. J Orthop Res. 2011;29(11):1775–1782.

17. Melvin A, Litsky A, Mayerson J, Witte D, Melvin D, Juncosa-Melvin N. An artificial tendon with durable muscle interface. J Orthop Res. 2010;28(2):218–224.

18. Abdullah S. Usage of synthetic tendons in tendon reconstruction. BMC Proceedings. 2015;9(3):A68.

19. Talia AJ, Tran P. Bilateral patellar tendon reconstruction using LARS ligaments: case report and review of the literature. BMC Musculoskeletal Disorders. 2016;17(1):302.

20. Hall PT, Stubbs C, Pedersen AP, Billings C, Stephenson SM, Greenacre CB, Anderson DE, Crouch DL. Effect of polyester-based artificial tendons on movement biomechanics: A preliminary in vivo study. Journal of Biomechanics. 2023;151:111520.

21. Barry JA, Cotter MA, Cameron NE, Pattullo MC. The effect of immobilization on the recovery of rabbit soleus muscle from tenotomy: modulation by chronic electrical stimulation. Exp Physiol. 1994;79(4):515–525.

22. Mathis A, Mamidanna P, Cury KM, Abe T, Murthy VN, Mathis MW, Bethge M. DeepLabCut: markerless pose estimation of user-defined body parts with deep learning. Nature Neuroscience. 2018;21(9):1281–1289.

23. Pataky TC. One-dimensional statistical parametric mapping in Python. Computer Methods in Biomechanics and Biomedical Engineering. 2012;15(3):295–301.

24. Pataky TC. Generalized n-dimensional biomechanical field analysis using statistical parametric mapping. J Biomech. 2010;43(10):1976–1982.

25. Pataky TC. 2022; https://spm1d.org. Accessed February 3, 2023.

26. Lieber RL, Blevins FT. Skeletal muscle architecture of the rabbit hindlimb: functional implications of muscle design. J Morphol. 1989;199(1):93–101.

27. Grover DM, Chen AA, Hazelwood SJ. Biomechanics of the rabbit knee and ankle: muscle, ligament, and joint contact force predictions. J Biomech. 2007;40(12):2816–2821.

28. Bojsen-Moller J, Magnusson SP. Mechanical properties, physiological behavior, and function of aponeurosis and tendon. J Appl Physiol (1985). 2019;126(6):1800–1807.

29. Bodine SC. Disuse-induced muscle wasting. Int J Biochem Cell Biol. 2013;45(10):2200–2208.

